# Intercellular adhesion molecule-1 protects against adipose tissue inflammation and insulin resistance but promotes liver inflammation and hepatic fibrosis in mice

**DOI:** 10.1101/2025.04.28.650968

**Authors:** Sreepradha Eswaran, Laura Gebert, Sarah Schraven, Nicole Treichel, Thomas Ritz, Sabine Hamm, Agnes Seeger, Fabian Kiessling, Thomas Clavel, Norbert Wagner, Angela Schippers

## Abstract

Metabolic dysfunction associated steatotic liver disease (MASLD) presents a growing global health problem with a range of manifestations, including steatosis, steatohepatitis, and cirrhosis. It is strongly associated with obesity, disease progression being promoted not only by hepatic leukocyte accumulation but also by inflammatory signals from adipose tissue and an altered gut microbiome. To determine the contribution of intercellular adhesion molecule-1 (ICAM-1) to MASLD pathogenesis, mice with an ICAM-1 mutation (Icam1^tmBay^) were compared to wild type (WT) mice in a Western-style diet (WD) model. WD-induced MASLD was accompanied by increased ICAM-1 expression in liver, epididymal white adipose tissue (EWAT), and intestine in WT mice. WD-fed Icam1^tmBay^ mice exhibited increased circulating neutrophils, higher frequencies of inflammatory leukocytes in EWAT, and a worsened glucose tolerance when compared to WT mice. In contrast, the mutation resulted in reduced WD-induced liver damage and less accumulation of intrahepatic leukocytes. WD-feeding caused substantial changes in fecal microbiota with decreased microbial diversity that differed between the mouse strains. In conclusion, ICAM-1 positively regulates adipose tissue homeostasis and protects from insulin resistance but promotes liver damage in diet-induced obesity. This points to various organ-specific roles for ICAM-1 and the potential of liver-specific targeting of ICAM-1 for treatment of MASLD.

## Introduction

Metabolic dysfunction-associated steatotic liver disease (MASLD) is strongly linked to obesity, hypertension, dyslipidemia, and insulin resistance and is considered to be the hepatic component of the metabolic syndrome ^1^. It represents a spectrum of liver diseases ranging from simple fat accumulation and metabolic dysfunction-associated steatohepatitis (MASH) to fibrosis and hepatocellular carcinoma. Steatosis-associated liver injury is now becoming recognized as the major cause of liver damage in Western societies ^2^. Inflammation of the liver is caused and maintained by an accumulation of leukocytes that are organized into inflammatory infiltrates, especially around damaged hepatocytes ^3,4^. This process is brought about by the interaction of the activated hepatic endothelium with chemokine receptors and cell adhesion molecules expressed on circulating immune cells ^5^.

The cell adhesion molecule intercellular adhesion molecule-1 (ICAM-1 or CD54) is a transmembrane glycoprotein member of the immunoglobulin (Ig) super-gene family expressed at a low level on leukocytes, endothelial-, epithelial-, and other cells ^6^. ICAM-1 expression is upregulated in response to inflammatory stimulation, particularly on endothelium, and best known for regulating the recruitment of circulating leukocytes to inflammatory sites. ICAM-1 binds to multiple ligands, including the β2 integrins LFA-1 (lymphocyte function-associated antigen-1, αLβ2), MAC-1 (macrophage-1 antigen, αMβ2), and p150,95 (αXβ2). Full-length membrane bound ICAM-1 consists of five extracellular Ig domains, a transmembrane domain, and a short cytoplasmic tail ^6^. The ligand LFA-1 binds to the first Ig domain, whereas Mac-1 and p150,95 have been shown to bind Ig domains 3 and 3/4, respectively. In addition to the full-length protein, six splice variants, varying in their combination and number of Ig domains, have been reported, which may also contribute to additional soluble (s) ICAM-1 variants. All the isoforms contain Ig domains 1 and 5. ^6,7^. Their expression seems to be cell-type specific and may depend on the inflammatory status and differentially impact disease outcome ^8^. Different results from experimental disease models using mice that lack different domains of the ICAM-1 protein support this hypothesis ^7,9^.

Leukocyte infiltration and low-grade inflammation are important characteristics of diet-induced metabolic dysfunction ^10^. ICAM-1 is important for leukocyte trafficking to inflammatory sites and contributes to the onset but also the resolution of pathogenesis ^6^. To address the impact of ICAM-1 in manifestations of metabolic dysfunction, we compared the outcome of western diet (WD)-induced MASLD in wild type (WT mice) and mice carrying a dysfunctional ICAM-1 gene (Icam1^tmBay^) ^9^. The Icam1^tmBay^ mice have a deletion of exon 5 which encodes the Ig domain 4 ^9^. They are not able to express the full-length protein, but due to alternative splicing they do express low amounts of the smaller ICAM-1 isoforms (isoforms containing domain 1+ 5, 1+2+5, and 1+2+3+5) ^7^. They show only residual membrane-bound ICAM-1 staining in thymus, lung, and spleen tissue but none at all in gut and liver ^11^. Here, we observed an increased inflammation of adipose tissue and a worsened glucose tolerance in WD-treated mice carrying the dysfunctional ICAM-1 gene. In contrast, WD-induced liver damage and intrahepatic immune cells accumulation were reduced in comparison to equally treated WT mice. In addition, WD-induced changes in fecal microbiota differed between the two mouse strains.

## Results

### Dysfunction of ICAM-1 has no impact on western diet-induced overall body fat deposition

Increased hepatic ICAM-staining and circulating ICAM-1 levels have previously been observed in MASH patients ^12^. In addition, elevated sICAM-1 levels have been detected in patients with metabolically unhealthy obesity and correlate with insulin resistance ^13,14^. Moreover, in high fat-treated mice, circulating sICAM-1 levels correlate with body and abdominal fat pad weight and male mice showed an upregulation of *Icam-1* mRNA in adipose tissue ^15^. To address the importance of ICAM-1 in manifestations of metabolic dysfunction, we analyzed the expression of ICAM-1 in experimentally induced MASLD. MASLD was induced by subjecting wild type (WT) mice for 24 weeks to a specific western diet (WD). Immunohistochemical expression of ICAM-1 was strongly increased in liver, epididymal white adipose tissue (EWAT), small intestine, and colon of WD-fed mice in comparison to the chow diet (CD)-fed controls (Fig. 1A). In addition, WD treatment caused significantly increased *Icam-1* mRNA levels in tissue homogenates of liver, EWAT and small intestine. (Fig. 1B-D). Next, we compared the outcome of WD-induced MASLD in WT mice and mice carrying the dysfunctional ICAM-1 gene (Icam1^tmBay^) ^9^. Weight gain in the ICAM-1 mutant was indistinguishable to that of WT mice (Fig. 1E) and both mouse strains showed the same cumulative food consumption (Fig. 1F). Furthermore, micro-computed tomography (µCT) after 12 weeks of WD feeding demonstrated similar body composition, quantified as body volume, liver volume, and visceral and subcutaneous fat volume (Fig. 1G-K), for both mouse strains.

**Figure 1.**
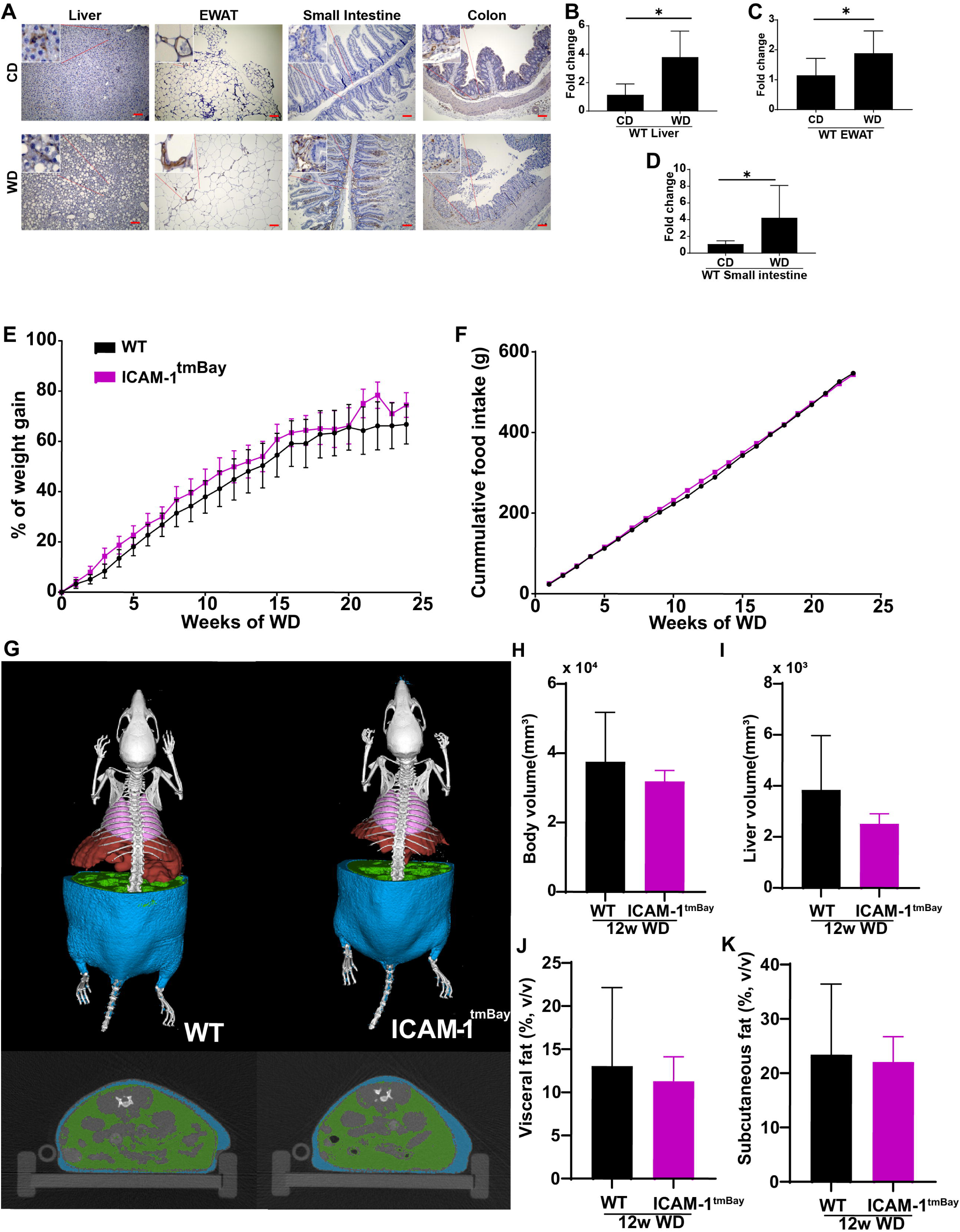
The ICAM-1 mutation has no impact on western diet-induced weight gain. Wild type (WT) mice (shown in black) and ICAM-1-mutant (Icam1^tmBay^) mice (shown in purple) were fed for 12 or 24 weeks with Western diet (WD). (**A**) Representative pictures of sections from the indicated organs (liver, epididymal white adipose tissue (EWAT), small intestine and colon) of WT mice that have been fed for 24 weeks with chow diet (CD) or WD. Sections have been stained with anti-ICAM-1 antibody and ICAM-1 protein is visualized as brown deposits developed with 3,3′-Diaminobenzidin reagent. Cellular nuclei are counterstained with hematoxylin (blue). Scale bar: 100 µm. (**B**-**D**) mRNA levels of *Icam1* were measured in (**B**) liver tissue (CD-fed (n = 3), WD-fed (n = 6) mice) (**C**) EWAT (CD-fed (n=11), WD-fed (n=12) and (**D**) small intestine tissue (CD-fed (n = 4), WD-fed (n = 7) of WT mice after 24 weeks of the indicated feeding. Values are expressed as fold increase over the mean value obtained for the respective CD-treated tissue. Statistical significance was calculated by non-parametric t-test. Values are represented as mean ± SD. * p ≤ 0.05. (**E**) Weight gain of WT (n = 8) and Icam1^tmBay^ (n = 8) mice represented as a percentage of weekly weight change, and (**F**) corresponding cumulative food intake. (**G**) Representative three-dimensional volume renderings of segmented bones (white), lungs (pink), visceral fat (green), and subcutaneous fat (blue) upon *in vivo* µCT imaging of 12 weeks WD-fed WT and Icam1^tmBay^ mice and 2D cross-sectional µCT images in transversal planes of the abdomen of the respective mice. **(H-K)** Quantification of the body composition of WT (n = 3) and Icam1^tmBay^ (n = 4) mice using µCT imaging after 12 weeks WD feeding **(G)** body volume, (**H**) liver volume, (**I**) visceral fat, and (**J**) subcutaneous fat. Statistical significance was calculated by *t*-test. Values are represented as mean ± SD.

### WD fed Icam1^tmBay^ mice display worsened glucose tolerance and inflammatory tone

Neutrophils are the first immune cells to accumulate in insulin-sensitive tissues during obesity and they seem to contribute to diet-induced peripheral tissue insulin resistance ^16^. Consistent with previous publications, the ICAM-1 mouse mutant displayed a higher percentage of circulating neutrophils when compared with WT mice (Fig. 2A) ^9^, which was further increased by treatment with WD. Increased circulating neutrophils in the ICAM-1 mutant did not, however, translate into a generally increased inflammatory activity of the myeloid blood cell population. Instead, the respective cell populations of both mouse strains displayed comparable phagocytic activity and ROS production (Supplementary Fig. S2). On regular CD, we found no difference in glucose homeostasis between WT mice and the ICAM-1 mutant. However, after 24 weeks of WD feeding Icam1^tmBay^ mice exhibited more pronounced features of the metabolic syndrome, as demonstrated by slower falls in glucose levels after glucose injection (Fig. 2B-C). Blood glucose levels of WD treated mice were independent of the genotype and slightly increased in comparison to CD-fed mice (Fig. 2 D) While serum triglyceride levels were unaffected by either diet or genotype, WD treatment caused increased levels of serum cholesterol, which did not vary between the mouse strains (Fig. 2E-F). Interestingly, TNFα and IL6 were significantly increased upon WD-feeding in Icam1^tmBay^ mice compared to WT mice. MCP-1 was slightly increased due to the diet in both mouse strains, but more in WT mice, and IL1B levels were not affected by diet or genotype (Fig. 2G.

**Figure 2.**
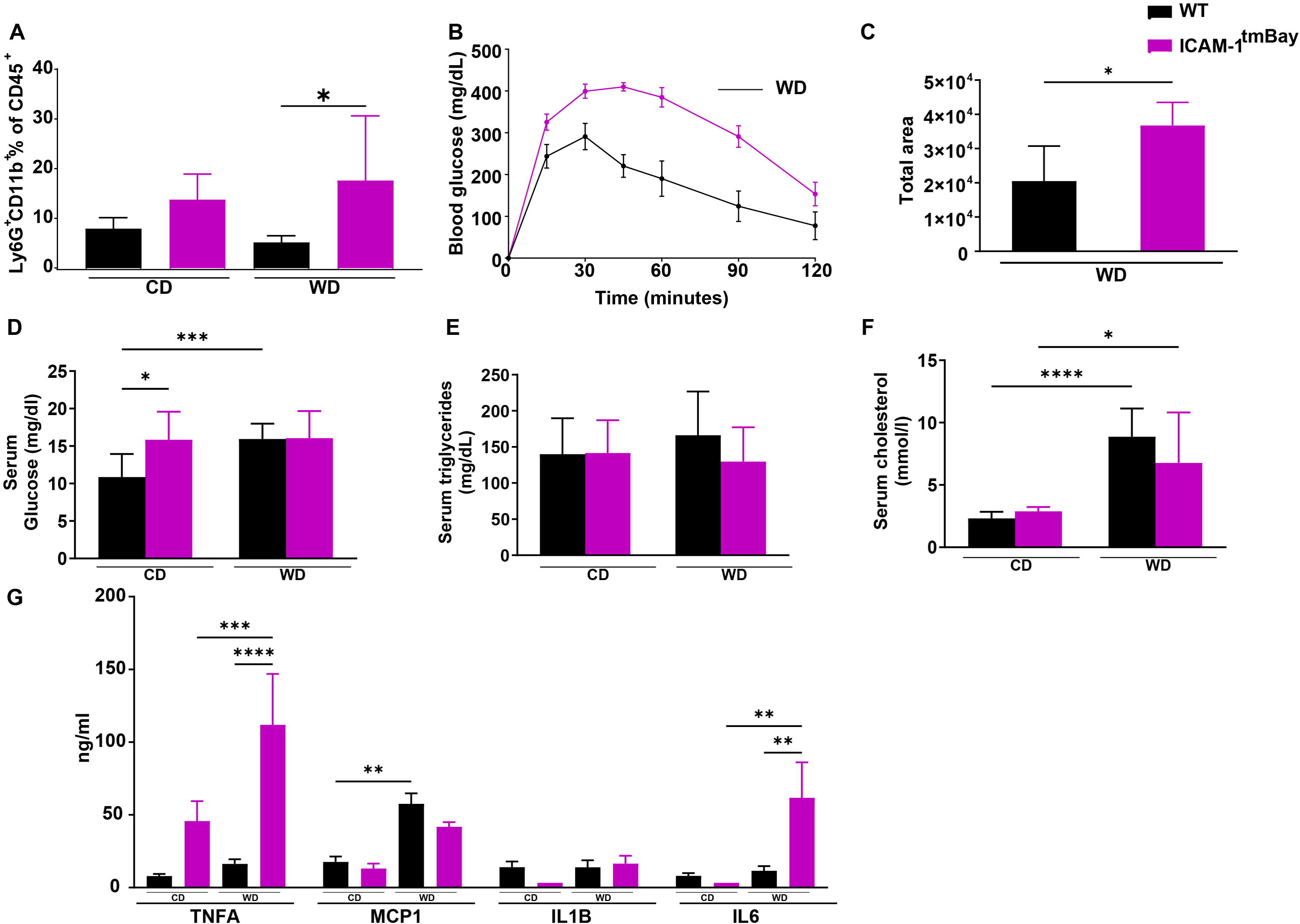
The ICAM-1 mutation aggravates glucose tolerance and systemic inflammation. Wild type (WT) mice (shown in black) and ICAM-1 mutant (Icam1tmBay) mice (shown in purple) were fed for 24 weeks with chow diet (CD, WT (n = 5-13), Icam1tmBay (n = 6)) or Western diet (WD, WT (n = 8-15), Icam1tmBay (n = 8)). (**A**) Comparative immune cell analysis of circulating neutrophils by flow cytometry. Representative FACS dot plots illustrating the gating strategy are shown in Supplementary Fig. S1. Depicted are the percentages of CD45^+^ cells of CD45^+^CD11b^+^Ly6G^+^ neutrophils. (**B**) Endpoint glucose tolerance test, and (**C**) corresponding quantification as area under curve. (**D**) Serum glucose, (**E**) serum triglycerides, and (**F**) serum cholesterol. (**G**) Serum pro-inflammatory cytokines tumor necrosis factor alpha (TNFA), monocyte chemoattractant protein-1 (MCP1), interleukin-1beta (IL1B), and interleukin-6 (IL6). Statistical significance was calculated either by the one-way ANOVA (**A-C**) or two-way ANOVA (**D-G**). Values are represented either as mean ± SD (**A**, **C-F**) or mean ± SEM **(B and G**). * p ≤ 0.05, ** p ≤ 0.01.

Together, these data demonstrate that upon WD-feeding Icam1^tmBay^ mice are more resistant glucose tolerance and display a higher inflammatory tone than respectively treated WT mice, however, no other metabolic feature was significantly altered due to the mutation.

### Increased accumulation of neutrophils and macrophages in adipose tissue of WD-fed Icam1^tmBay^ mice

MASLD is strongly associated with obesity, and inflamed adipose tissue seems to promote the progression to MASH ^4,10^. In lean mice and humans, macrophages constitute 5% of cells in adipose tissue depots, but in obese rodents and humans macrophages are increased by up to 50% ^17^. In both mouse strains, WD feeding caused an increased EWAT to body weight ratio when compared to respective CD-fed mice, with no difference between the two mouse strains (Fig. 3A). Histological staining of macrophages in EWAT using an antibody against CD68^+^ showed the arrangement of macrophages in characteristic crown-like structures (CLS) in both groups of obese WD-fed mice but not in lean CD-fed mice (Fig. 3B, CLS are marked with black arrows). CLS around dead adipocytes are indicators of pathological changes in fatty tissue ^18^ and positively correlate with metabolic disorders. CLS in adipose tissue of the WD-fed ICAM-1 mutant were slightly increased in comparison to comparably treated WT mice. This was supported by flow cytometric analysis where we observed significantly increased frequencies of CD11b^+^ Ly6G^+^ neutrophils and CD11b^+^ F4/80^+^ macrophages in adipose tissue of the WD-treated Icam1^tmBay^ mice (Fig. 3C-D). The frequencies of CD4^+^ or CD8^+^ T cells remained unchanged by diet and genotype (not shown).

**Figure 3.**
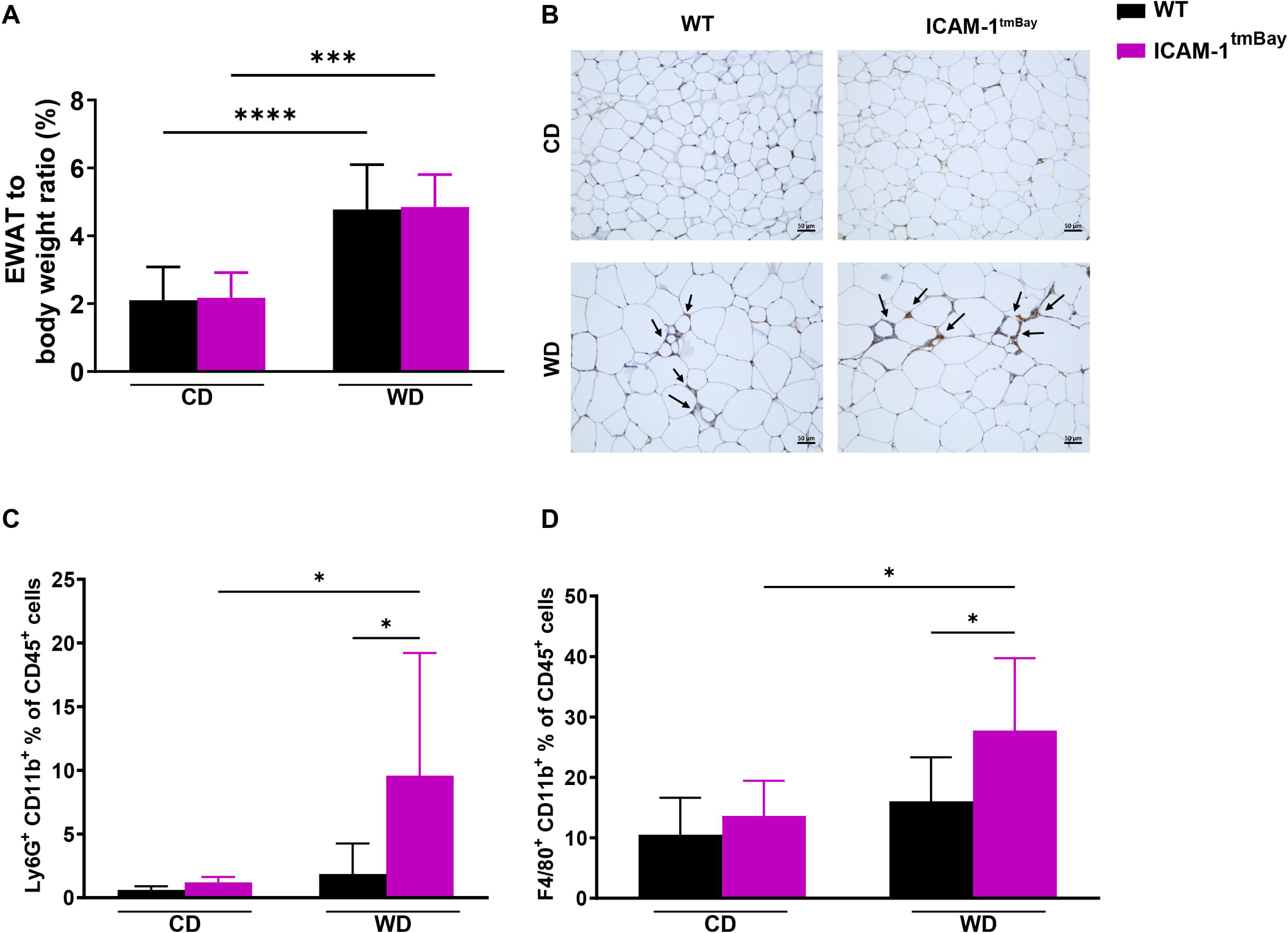
Epidydimal white adipose tissue of WD-fed Icam1^tmBay^ mice exhibits increased inflammation of immune cells. Wild type (WT) mice (shown in black) and Icam1^tmBay^ mice (shown in purple) were fed for 24 weeks with chow diet (CD) or Western diet (WD). Depicted are the endpoint values of WT CD-fed (n=6-13), WT WD-fed (n=8-15), Icam1^tmBay^ CD-fed (n=6), and Icam1^tmBay^ WD-fed (n=8). (**A**) EWAT/body weight ratio. (**B**) Representative images of CD68 stained Epidydimal white adipose tissue (EWAT) sections of the indicated mice strains (original magnification X 20, scale bar = 50 µm). Areas with CD68^+^ crown-like structures are labeled with black arrows. (**C-D**) Comparative immune cell analysis of EWAT by flow cytometry. Representative FACS dot plots illustrating the gating strategy are shown in Supplementary Fig. S1. Depicted are the percentages of CD45^+^ cells of (**C**) neutrophils (CD45^+^CD11b^+^F4/80^-^Ly6G^+^), and (**D**) macrophages (CD45^+^Ly6G^-^CD11b^+^F4/80^+^). Statistical significance was calculated by the two-way ANOVA. Values are represented as mean ± SD (**A**, **C-D**). * p ≤ 0.05, *** p ≤ 0.001, **** p ≤ 0.0001.

### ICAM-1 Dysfunction Ameliorates MASH Progression

Under basal conditions of chow feeding, WT mice and mice with the dysfunctional ICAM-1 gene displayed no signs of hepatocyte injury, as manifested in low serum values of AST, ALT, and ALP (Fig. 4A-C). However, WD feeding led to significantly increased levels of serum transaminases and ALP in WT mice, which was not observed in the respectively treated ICAM-1 mutant. This may suggest that the ICAM-1 mutation protects mice to some extent from liver damage.

**Figure 4.**
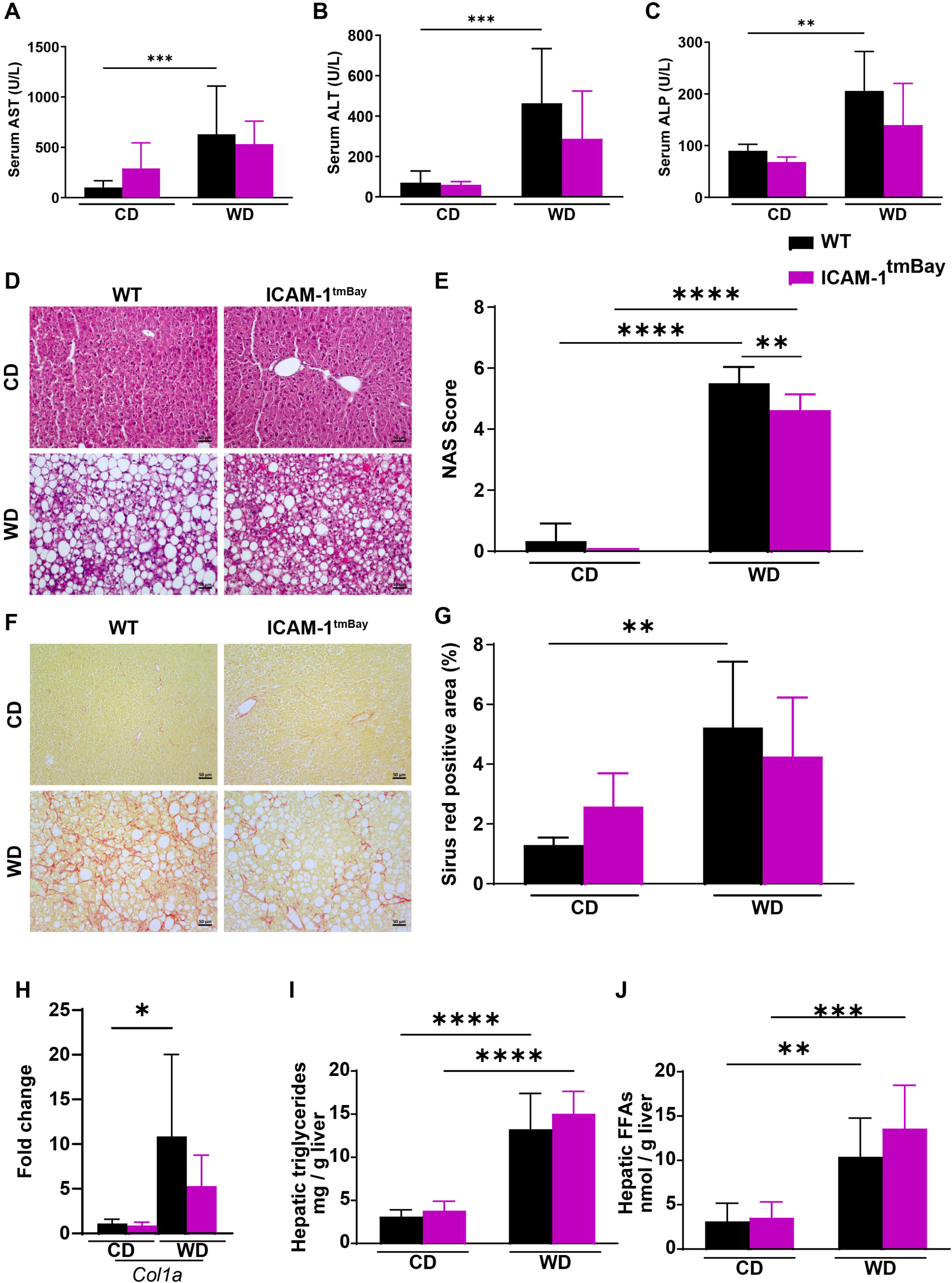
WD-fed Icam1^tmBay^ mice exhibit ameliorated MASH progression. Wild type (WT) mice (shown in black) and Icam1^tmBay^ mice (shown in purple) were fed for 24 weeks with chow diet (CD) or Western diet (WD). Depicted are the endpoint values of WT CD-fed (n=4-13), WT WD-fed (n=7-15), Icam1^tmBay^ CD-fed (n=6), and Icam1^tmBay^ WD-fed (n=6-8). (**A**) Quantification of serum aspartate aminotransferase (AST). (**B**) Quantification of serum alanine aminotransferase (ALT). (**C**) Quantification of alkaline phosphatase (ALP). (**D**) Representative images of H&E-stained liver sections of the indicated mice strains. Black arrows indicate the presence of inflammatory foci (original magnification X 20, scale bar = 50 µm). (**E**) Non-alcoholic fatty liver disease (NAFLD) activity score (NAS). (**F**) Representative images of Sirius red-stained liver sections of the indicated mice strains (original magnification X 20, scale bar = 50 µm). (**G**) Quantification of Sirius red-positive area expressed as a percentage of the tissue area. (**H**) mRNA levels of collagen 1alpha (*col1a*) in liver tissue homogenates. For quantification, values are expressed as fold increase over the mean values obtained for liver tissue from CD-fed WT mice. (**I**) Hepatic triglycerides in liver tissue homogenates. (**J**) Free fatty acids (FFA) in liver tissue homogenates. Statistical significance was calculated by the two-way ANOVA. Values are represented as mean ± SD (**A-C**, **E**, **G**, **H**). * p ≤ 0.05, ** p ≤ 0.01, *** p ≤ 0.001, **** p ≤ 0.0001.

Histological evaluation of H&E-stained liver sections revealed a marked increase in hepatocyte ballooning with micro- and macrosteatosis in both mouse strains after 24 weeks of WD (Fig. 4D). However, this was significantly less pronounced in the Icam1^tmBay^ mice, as can be seen in the respective NAFLD activity score (NAS) (Fig. 4E). Moreover, Sirius red staining of liver sections revealed a marked increase in hepatic fibrosis in WT mice post WD feeding, whereas comparably treated mutant mice were less affected (Fig. 4F-G). In line with our observations of an increased WD-induced fibrosis, both mouse strains showed increased *Col1a* mRNA post WD feeding in liver tissue homogenates. This increase was less pronounced in the ICAM-1 mutant (Fig. 4H). No differences were observed in levels of hepatic triglycerides or free-fatty acids between the four experimental mice groups (Fig. 4I, J). Increased intrahepatic inflammatory immune cell proportions are a characteristic feature of progressing MASLD. Therefore, we next performed a detailed flow cytometric analysis of the major hepatic leukocyte groups. (Fig. 5A). Feeding with WD caused an overall increase in the frequencies of CD4^+^ T cells and macrophages, that was independent of the genotype, whereas the frequency of B cells was decreased. Under chow diet conditions, we observed lower proportions of hepatic CD8^+^ T cells in the ICAM-1 mutant. However, 24 weeks post WD feeding, both mouse strains showed the same frequencies of hepatic CD8^+^ T cells. The WD-induced increase in macrophages, which are central players in MASLD progression, was less pronounced in Icam1^tmBay^ mice. This result is in line with a decreased expression of *Mcp-1* mRNA as detected by RT-PCR on liver tissue homogenates. (Fig. 5B).

**Figure 5.**
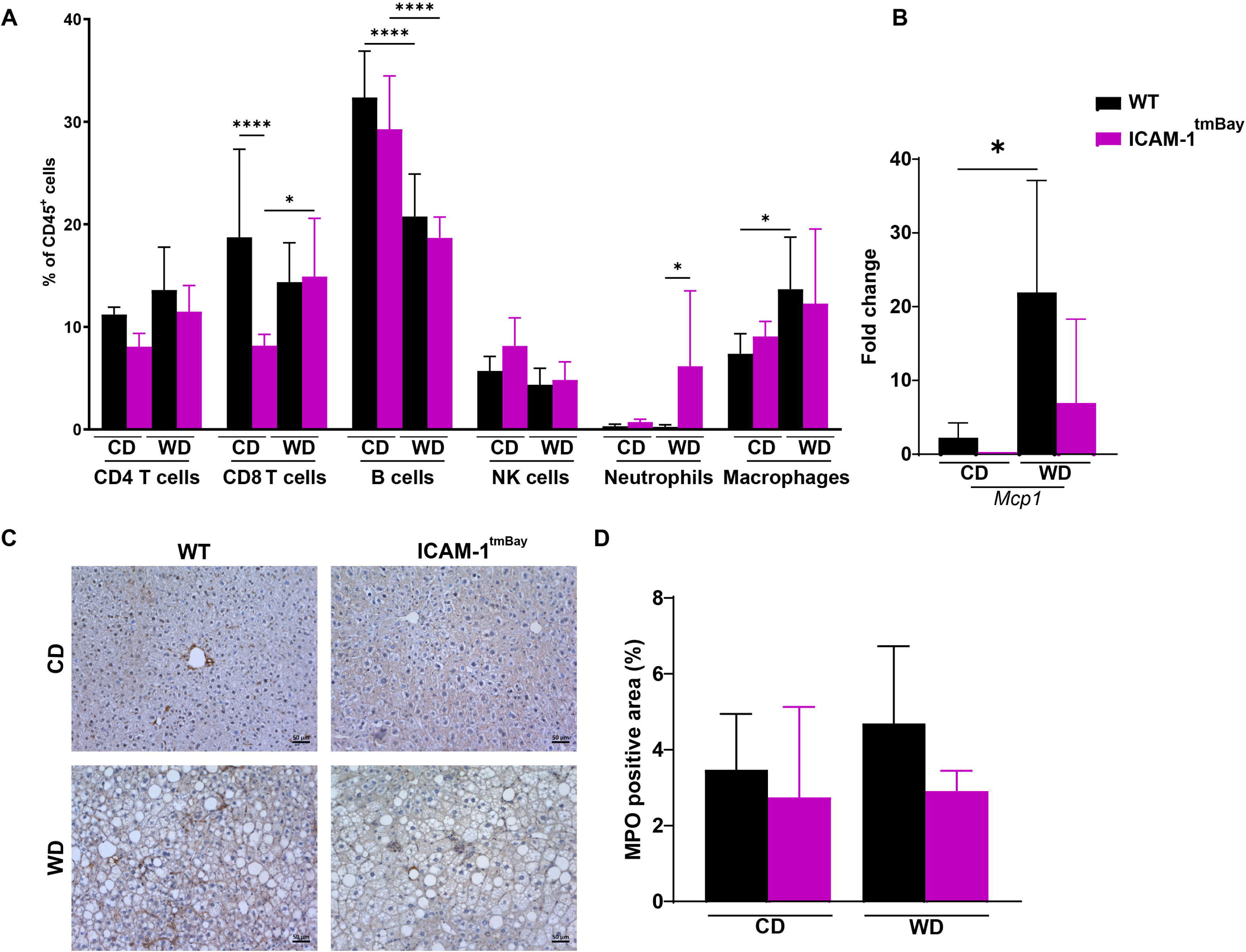
Increased WD-induced hepatic neutrophil frequencies in Icam1^tmBay^ mice do not correlate with hepatic myeloperoxidase expression. Wild type (WT) mice (shown in black) and mice with dysfunctional ICAM-1 gene (Icam1^tmBay^) mice (shown in purple) were fed for 24 weeks with chow diet (CD) or Western diet (WD). Depicted are the endpoint values of WT CD-fed (n=4-6), WT WD-fed (n=7-8), Icam1^tmBay^CD-fed (n=5-6), and Icam1^tmBay^WD-fed (n=7-8). (**A**) Comparative immune cell analysis of liver by flow cytometry. Representative FACS dot plots illustrating the gating strategy are shown in Supplementary Fig. S1. Depicted are the percentages of CD45^+^ cells of CD4^+^ T cells (CD45^+^CD3^+^CD8^-^CD4^+^), CD8^+^ T cells (CD45^+^CD3^+^CD4^-^CD8^+^), B cells (CD45^+^CD3^-^CD19^+^), NK cells (CD45^+^NK1.1^+^), neutrophils (CD45^+^CD11b^+^Ly6G^+^), and macrophages (CD45^+^Ly6G^-^CD11b^+^F4/80^+^). (**B**) mRNA levels of monocyte chemoattractant protein 1 (*Mcp1*) in liver tissue homogenates. For quantification, values are expressed as fold increase over the mean values obtained for liver tissue from CD-fed WT mice. (**C**) Representative images of myeloperoxidase (MPO) stained liver sections of the indicated mice strains (original magnification X 20, scale bar = 50 µm). (**D**) Quantification of MPO positive area expressed as a percentage of the tissue area. Statistical significance was calculated by two-way ANOVA. Values are represented as mean ± SD (**B**, **C**). * p ≤ 0.05, ** p ≤ 0.01, **** p ≤ 0.0001.

Interestingly, a WD induced increase in hepatic neutrophil frequencies, was only observed in the ICAM-1 mutant. Expression of Myeloperoxidase (MPO), an enzyme predominantly released by neutrophils, correlates with the presence of MASH in patients and murine livers fibrosis, and MPO deficiency or pharmacological inhibition protects against MASH-induced liver injuries in mice ^19^. To assess neutrophil-mediated liver damage, we performed immunohistochemical staining for MPO on liver sections (Fig. 5C). Most interestingly, the increased WD-induced neutrophil frequencies in Icam1^tmBay^ mice did not result in an increased MPO signal intensity. In fact, the MPO positive area in liver sections from the WD-treated ICAM mutant was smaller than the area in respectively treated WT mice (Fig. 5D). Although this finding was not statistically significant, it correlated with the overall MASLD amelioration observed in mice with dysfunctional ICAM-1 expression.

### Fecal microbiota profiles are altered by WD-feeding

Low-grade inflammation triggered by WD is attributed in part to an increased translocation of bacterial products into the circulation, involving an altered microbiome and increased intestinal permeability ^20^. To compare WD-induced changes in the gut microbiota of WT and ICAM-1^tmbay^ mice, we used 16S rRNA gene amplicon sequencing. Fecal samples were collected from WD-fed WT and ICAM-1^tmbay^ mice at the start (0 weeks, 16 samples) and end (24 weeks, 14 samples) of the experimental feeding. Across the 30 samples sequenced, 897,246 high-quality, chimera-checked sequences (29,908 ± 15,279 per sample) were analyzed, representing 222 operational taxonomic units (OTUs) (135 ± 69 OTUs per sample). Sufficient sequencing depth for all samples was estimated based on the plateauing of the rarefaction curves (Supplemental Fig. S3). First, we assessed differences in bacterial profiles between the experimental groups at the start and end point of WD-treatment by meta non-metric multidimensional scaling (NMDS) visualization of Unifrac distances (Fig. 6A). WD feeding led to significantly altered microbiota profiles in both mouse strains. Alpha-diversity was then evaluated by calculating bacterial richness (i.e., number of observed OTUs) (Fig. 6B) and Shannon effective counts (Fig. 6C). At the starting point a higher diversity was observed in ICAM-1^tmBay^ mutant mice, which decreased significantly after 24 weeks of WD feeding. Next, the relative abundance of major bacterial phyla was calculated (Fig. 6D-G). Whilst the proportions of Bacillota (formerly Firmicutes), Actinomycetota (formerly Actinobacteria) and Pseudomonadota (formerly Proteobacteria) were significantly increased and that of Bacteroidota (formerly Bacteroidetes) significantly decreased in feces of WD-fed WT mice, their relative abundance seemed to be unaffected by WD feeding in feces of the ICAM-1^tmBay^ mutant mice. Both mouse strains exhibited equal relative abundances of Bacillota and Bacteroidota after WD feeding. In contrast, the relative abundances of Actinomycetota and Pseudomonadota were substantially higher in feces of the WT mice. The most obvious WD-induced differences at the family level (Figure 7H-L) were an increase in the relative abundance of *Bacteroidaceae* and *Peptostreptococcaceae* and a decrease in the relative abundance of *Muribaculaceaea* which was unaffected by the mouse genotype. Interestingly, increased proportions of *Clostridiaceae* were only observed in WT mice, while these were practically absent in feces of ICAM-1^tmBay^ mutants. In contrast, an increased relative abundance of *Prevotellaceae*, which was absent in the WT, was only seen in the feces of ICAM-1^tmBay^ mutants.

**Figure 6.**
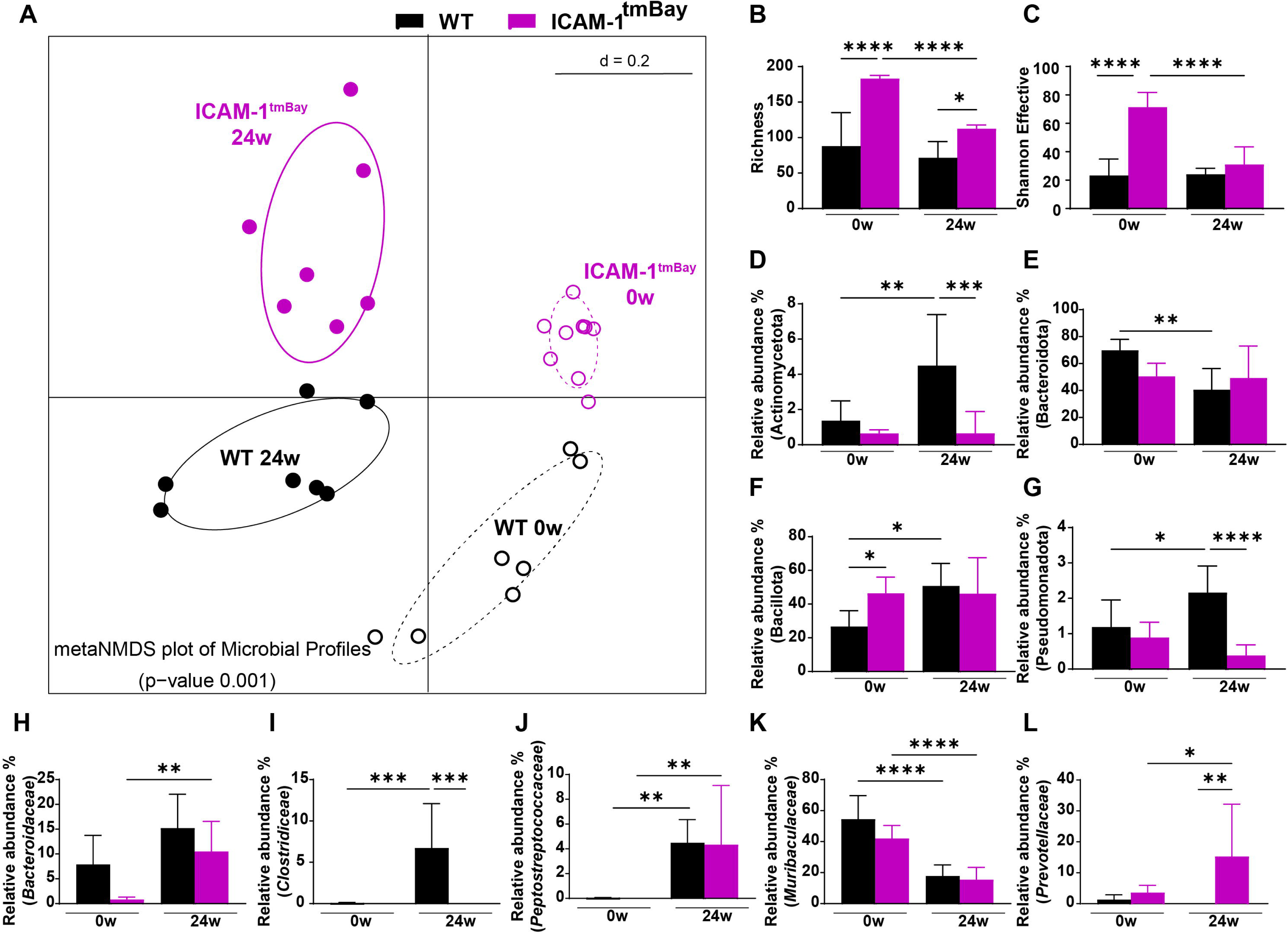
WD feeding causes important shifts in gut microbiota diversity and composition. Fecal samples were collected from wild type (WT) mice (black) and mice with dysfunctional ICAM-1 gene (Icam1^tmBay^) (purple) before (0 weeks (0w)) and after 24 weeks (24w) of western diet (WD) feeding. Number of mice: WT 0w (n=7), WT 24w (n=7), Icam1^tmBay^ 0w (n=9), and Icam1^tmBay^ 24w (n=7). The diversity of operational taxonomic units (OTUs) within a given sample (alpha-diversity) and between samples (beta-diversity) was calculated in R using Rhea ^51^. (**A**) Beta-diversity of experimental groups shown as meta non-metric multidimensional scaling (NMDS) plot of microbiota profiles based on generalized UniFrac distances from mice at the indicated time points. Individual time points that were considered for calculations have been marked within their respective circles. Alpha-diversity of experimental groups shown as (**B**) number of observed OTUs (richness), or (**C**) Shannon effective index. (**D**-**G**). Relative abundance at the phylum level. (**H**-**L**) Relative abundance at the family level. Statistical significance was calculated by PERMANOVA (**A**) or two-way ANOVA (**B-L**). Values are represented as mean ± SD (**B-L**). * p ≤ 0.05, ** p ≤ 0.01, ***p ≤ 0.001, **** p ≤ 0.0001.

## Discussion

MASLD progression involves a complex interplay of imbalanced inflammatory cell populations and inflammatory signals derived from several organs including liver, adipose tissue, gut and its enteric microbiota, making MASLD rather to a systemic disease ^1^. The progression from steatosis to cirrhosis is driven by chronic inflammation of the injured liver, involving adhesion molecule-mediated immigration of inflammatory immune cells into the liver parenchyma ^3^. However, these processes have not yet been sufficiently elucidated, which is prerequisite for identifying therapeutic targets.

We set up experiments to better define the role of the adhesion molecule ICAM-1 in WD-induced MASLD. Previously, increased ICAM-1 expression in adipose tissue has been shown in mice fed for 2, 3 or 6 months on a high-fat or high-fat/high-sugar diet ^15,21^. Here we show that WD treatment also leads to increased ICAM-1 expression in liver, adipose tissue,colon, and small intestine of WT mice, pointing to an obesity-induced inflammatory status within those tissues and supporting the notion of their joint involvement in MASLD progression. ICAM-1 does not appear to be involved in WD-induced food intake and body fat gain, as there were no differences in overall body fat deposition between WD-fed WT mice and Icam1^tmBay^ mice carrying the dysfunctional ICAM-1 gene. These observations corroborate the results of earlier studies where no differences in the weight of fat pads were detected in WD-fed WT-mice and ICAM-1^null^-mice ^22^. Moreover, mice deficient for the ICAM-1 ligand Mac-1 and mice lacking β2 integrin function exhibited identical WD-induced weight gain and expanded fat mass ^22,23^. WD-fed Icam1^tmBay^ mice exhibited an increased insulin resistance and systemic inflammation, as evidenced by worsened glucose tolerance and increased circulating neutrophils and TNFα, suggesting a protective effect of ICAM-1 against metabolic dysregulation. In contrast, an overall improvement in glucose tolerance has been observed in WD-treated mice lacking beta2-integrin function ^23^, suggesting that other ICAM-1 ligands may counteract the negative impact of beta2-integrin ligands on glucose metabolism. Systemic glucose regulation is complex and involves several organs, including adipose tissue, where inflammatory status directly impacts on insulin sensitivity. Obesity is accompanied by the progressive recruitment of pro-inflammatory immune cells to adipose tissue, which contribute to the development of metabolic dysregulation ^24^. Inflammatory macrophages, which differentiate from monocytes accumulate in adipose tissue and are thought to play a key role in adipose tissue inflammation ^17^. The higher frequencies of macrophages in the adipose tissue of ICAM-1 mutants observed here could therefore be causally involved in the impaired glucose tolerance.

In contrast, to the worsened WD-induced metabolic phenotype upon ICAM-1 dysfunction, we observed a slower MASH progression in Icam1^tmBay^ mice, reflected in lower serum liver damage levels, a lower NAS score and reduced fibrosis compared to respectively treated WT animals, suggesting that ICAM-1 contributes to disease progression. Amelioration of disease was accompanied by a smaller increase in the frequency of hepatic macrophages and decreased expression of MCP-1, the chemokine ligand known to be responsible for augmented infiltration of inflammatory monocytes ^25^. As bone marrow-derived hepatic macrophages reportedly display an inflammatory phenotype and contribute to hepatocellular damage in experimental models of chronic liver injury and obesity ^26,27^, it would be reasonable to attribute reduced inflammatory macrophages in livers of Icam1^tmBay^ mice to the observed amelioration of disease. In addition to the finding of increased circulating neutrophils in the ICAM-1 mutant in homeostasis, which corroborates earlier results ^9^, we observed higher percentages of neutrophils in adipose tissue and liver tissue of the WD-fed mutant, a phenotype which has also been seen in mice lacking beta2-integrin function ^23^. We thereby conclude that WD-induced neutrophil migration to liver and fat can also occur independently of ICAM-1 and that increased neutrophil numbers in these tissues might result from the strongly increased levels of circulating neutrophils. This hypothesis fits with previously published data showing that acute LPS-induced neutrophil infiltration in the cremaster muscle and peritonitis model does not require ICAM-1 ^28^. Interestingly, higher frequencies of neutrophils in WD-fed Icam1^tmBay^ mice did not translate into an increased inflammatory activity of these cells, as shown by a rather lower MPO expression in liver tissue in comparison to the WT situation. This is in line with the improved liver phenotype. The finding points to a reduced activation potential of neutrophils with dysfunctional ICAM-1, which would also explain why phagocytic capacity and ROS production of leukocytes in blood from mutant mice remained stable despite increased numbers of circulating neutrophils. Of note, previously it was shown that LPS induced neutrophil ICAM-1 expression and ICAM-1-mediated intracellular signaling caused increased phagocytosis and ROS generation ^28^.

Diet-induced gut microbiota changes and a leaky barrier contribute to the systemic inflammation and disease progression of MASLD ^20^. In addition to alterations in gut permeability, WD-induced gut microbiota changes can affect hepatic lipid and carbohydrate metabolism, inflammatory processes in liver and adipose tissue through activation of innate immunity, gut hormone production, and bile acid metabolism ^29^. Moreover, a contribution of the enteric microflora to the constitutive expression of ICAM-1 in intestinal and liver tissue has been reported ^30^. In future, gut microbiota manipulations could become a useful tool for the treatment of obesity-related diseases while potential influences of the gut microbiota on ICAM-1-expression and vice versa could be of interest. At the family level, fecal microbial composition of both mouse strains exhibited a substantial increase in the relative abundance of *Bacteroidaceae* and *Peptostreptococcaceae* after 24 weeks of WD. Increased relative abundance of *Peptostreptococcaceae* (phylum Bacillota) after WD feeding has already been demonstrated in previous studies and might present a marker of disease risk ^31^. According to Iwamoto et al., *Peptostreptococcaceae* contribute to cholesterol conversion and might contribute to WD-induced hypercholesterolemia ^32^. The WD-induced increase in relative abundance of *Bacteroidaceae* (phylum Bacteroidota) observed here also fits with previous publications reporting increased *Bacteroidaceae* in high-fat diet treated mice and a positive correlation of the genus *Bacteroides* (family *Bacteroidaceae*) with disease severity in patients with biopsy proven NASH ^33,34^. *Bacteroides*-associated hepatic pathology is associated with changes in bacterial metabolites such as a decrease in short chain fatty acids or increases in deoxycholic acid and carbohydrates which can promote liver inflammation ^35^. Independent of the genetic background, WD caused a drastic decrease in the *Muribaculaceae*, which represent a large proportion of the mouse gut Bacteroidota. This corroborates former studies showing a decreased relative abundance of *Muribaculaceae* upon WD feeding and a negative correlation with obesity ^36^. Interestingly, a positive correlation of these bacteria with life span has been observed in mice. Furthermore, feces of WD-fed WT mice, but not of ICAM-1^tmbay^ mutant mice, showed a significantly higher relative abundance of *Clostridiaceae*. A WD-induced expansion of *Clostridaceae* (phylum Bacillota) has been reported previously ^37^. *Clostridium* species contribute to bile acid metabolism and, according to Xiao et al. ^38^, expansion of these bacteria might contribute to an altered bile acid metabolism upon WD feeding. Unexpectedly, we observed an increased relative abundance of *Prevotellaceae* (phylum Bacteroidota) in feces of WD fed ICAM-1^tmbay^ mice, which was not seen in WT mice. According to the literature, the occurrence of *Prevotellaceae* is negatively related to a high-fat diet in mice, pigs and humans ^39^. In addition*, Segatella copri* (formerly *Prevotella copri*) ^40^ has been reported to improve the abnormal glucose metabolism in diabetic mice and ameliorated histological damage of pancreas, liver and colon ^41^. An increased relative abundance in the *Prevotellaceae* might therefore have contributed to improvement of liver damage in the ICAM-1^tmbay^ mutant. Collectively, our study reveals marked changes in the gut microbiota of WT and ICAM-1^tmbay^ mice due to WD feeding, and differences in the microbiota between both mouse strains, which might have contributed to the different phenotypes.

In summary, we have shown that ICAM-1 has no impact on WD-induced body fat gain, protects from systemic glucose tolerance and differentially alters local tissue inflammation by ameliorating adipose tissue inflammation and aggravating MASH progression. This points to a variety of organ-specific roles for ICAM-1 and the potential of liver-specific targeting of ICAM-1 for treatment of MASLD.

## Methods

### Housing, Mice, and Dietary Treatments

Animals were housed in specific pathogen-free conditions with 12 h light/dark cycles and water and food available ad libitum. The study was carried out in accordance with ARRIVE guidelines and the regulations laid down by the regional authorities for nature, environmental, and consumer protection of North Rhine-Westphalia (LANUV, Recklinghausen, Germany) and approved by the respective Committee (Permit Number: 81-02.04.2017.A429). Western diet (WD) treatments were performed with 8 to 12-week-old male only, as female mice are more resistant to diet-induced metabolic disorders ^42^. ICAM-1 mutant (Icam1^tm1Bay^) mice ^9^ and wild type (WT) mice on C57BL/6J background from the same barrier and room weighing at least 25 g were fed chow diet (CD) (9 kcal % fat, 24 kcal % protein, 67 kcal % carbohydrates) (ssniff, Soest, Germany, rat/mouse–maintenance) or WD (40 kcal % fat (vegetable fats, 20 kcal % fructose, 2% cholesterol) (Brogaarden, Lynge, Denmark; cat. no. D16022301). All WD experiments were repeated in at least two independent experimental setups.

### Serum and Liver Biochemical Measurements

Serum aspartate aminotransferase (AST), serum alanine aminotransferase (ALT), alkaline phosphatase (ALP), glucose, triglyceride, and cholesterol levels in serum were measured by the Central Laboratory Facility of the University Hospital, RWTH Aachen.

Determination of the intrahepatic triglyceride concentration was performed in accordance with the manufacturer’s instructions of the Instruchemie LiquiColor mono Kit (Instruchemie, Delfzijl, the Netherlands). Intrahepatic free fatty acid quantification was measured with the FFA quantification kit (Abcam, Cambridge, UK, cat. no ab65345) as instructed by the manufacturer. Serum cytokine concentrations were measured by a bead-based immunoassay technique using the LegendPlex Mouse Inflammation Panel (13-plex) (Biolegend, San Diego, cat. no 740446) following the manufacturer’s instructions. Samples were acquired on a Canto-II flow cytometer (BD Biosciences) and data were analyzed using the LegendPlex data analysis software.

### Glucose Tolerance Test (GTT)

Mice were fasted for 6 h and blood glucose was measured using an Accu-Check^®^ Aviva meter (Roche, Basel, Switzerland), via one drop of blood taken from the animal’s tail, every 15 min for 2 h following intraperitoneal administration of 2 g/kg glucose.

### Micro-Computed Tomography (μCT)

In vivo μCT imaging was performed using a hybrid µCT-Fluorescence Tomography system (MILabs B.V., Houten, the Netherlands) in the ultra-focus fast scan mode. The X-ray tubes of the μCT were operated at a voltage of 65 kV with a current of 0.13 mA. To cover the entire mouse, a continuous rotation scan was performed with one full rotation, exposure time of 75 ms, total scan duration of 27s, and dose estimation of 35 mGy. After acquisition, volumetric data sets were reconstructed at an isotropic voxel size of 140 μm using MILabs Auto Rec 12.00. The fat-containing tissue regions, which appear hypo-intense in the μCT data, were segmented using an automated segmentation method with interactive correction of segmentation errors using the Imalytics Preclinical 3.0.2.5 software (Gremse-IT GmbH, Aachen, Germany) ^43^.

### Histological Stainings

All stainings were performed on 4 μm paraffin sections. Images were acquired using an Axioplan2 microscope (Carl Zeiss Microscopy, Oberkochen, Germany) and Zen lite 3.2 software (Carl Zeiss Microscopy, Oberkochen, Germany). Histopathological scoring of hematoxylin and eosin stained sections from liver and their validation was performed blinded via a non-alcoholic fatty liver disease (NAFLD) activity score (NAS) ^44^. The total hepatic parenchymal area and the Sirius red positive area were estimated by means of a size marker using ImageJ software (version 1.50; National Institutes of Health Bethesda, MD ^45^).

Immunohistochemical stainings were performed with a rabbit-anti-ICAM-1, –MPO, or –CD68 antibody and biotinylated goat-anti-rabbit IgG. Antibody binding was visualized with the ABC kit and diaminobenzidine reagent as a chromogen (both from Vector, Burlingame, USA). Antibodies are listed in the Supplementary materials Table S1.

### Flow Cytometry

All flow cytometric measurements were performed on a Canto-II cytometer (BD Biosciences). Single cell suspensions were stained directly using combinations of the monoclonal antibodies listed in Supplementary materials Table S2. Representative FACS dot plots illustrating the gating strategy are shown in Supplementary Fig. S1. Data were analyzed by FlowJo 10.2 software (Tree Star, Ashland, OR, USA).

### Gene Expression Analysis by Real-Time PCR

Real-time polymerase chain reactions (RT-PCR) were performed in duplicate on a 7300 RT-PCR system with 7000 System SDS Software Version 1.2.3 (Applied Bioscience, Darmstadt, Germany) using the quantitative (q)PCR Master Mix for SYBR Green I (Eurogentec, Cologne, Germany). Glyceraldehyde 3-phosphate dehydrogenase (GAPDH was used as endogenous control for normalization). Primer sequences are listed in Supplementary materials Table S3.

### DNA Isolation from Feces, 16S rRNA Gene Amplicon Sequencing, and Data Analysis

Stool samples were processed by the Functional Microbiome Research Group at the Faculty of Medicine, RWTH Aachen University. Metagenomic DNA was isolated from samples and negative controls using a modified protocol according to Godon et al.^46^ 16S rRNA gene amplicons were generated with primers 341F-785R ^47^ and a two-step PCR (15 + 10 cycles) ^48^ using a pipetting robot. Sequencing was performed as described in detail previously ^49^ Individual libraries were diluted to a final concentration of 4 nM and 5 µl of each library were added to the final pool. The 16S rRNA gene amplicon libraries were sequenced with dual barcodes in paired-end mode (2 x 300 nt) using a MiSeq sequencer (Illumina, Inc.). Data were analyzed with an updated version of a previously described workflow ^49^. Raw reads were processed using an in-house developed pipeline (www.imngs.org) ^50,51^ based on UPARSE. In brief, sequences were demultiplexed and trimmed to the first base with a quality score < 3. The pairing, chimaera filtering and OTU clustering (97 % identity) was done using USEARCH 11.0 ^52^. Sequences with less than 350 and more than 500 nucleotides and paired reads with an expected error >3 were excluded from the analysis. The remaining reads were trimmed by ten nucleotides on each end to avoid GC bias and non-random base composition. Operational taxonomic units (OTUs) were clustered at 97 % sequence similarity and only those with a relative abundance > 0.25 % in at least one sample were kept. Sequence alignment and taxonomic classification were conducted with SINA 1.6.1, using the taxonomy of SILVA release 128 ^53^. Downstream analysis was performed in the R programming environment using Rhea (https://lagkouvardos.github.io/Rhea/) ^51^. OTU tables were normalized to account for differences in sequence depth. Βeta-diversity was computed based on generalized UniFrac distances ^54^. Αlpha-diversity was assessed based on species richness and Shannon effective diversity ^55^.

### Statistical Analysis

Unless otherwise indicated, statistical analysis was performed with GraphPad Prism software (version 10 GraphPad, La Jolla, CA, USA). Data are presented as mean ± Standard Deviation (SD). Significance values were calculated using the Student’s t-test when comparing two groups or two-way analysis of variance (ANOVA) and Tukey post-test. Values of p < 0.05 were considered significant (* p < 0.05, ** p < 0.01, *** p < 0.001, and **** p ≤ 0.0001).

## Acknowledgments

This research was funded by the DFG (German Research Foundation)–Project-ID 403224013– SFB 1382 to T.C., N.W., F.K., and A.S. This work was supported by the “Immunohistochemistry facility”, a core facility of the Interdisciplinary Center for Clinical Research (IZKF) Aachen within the Faculty of Medicine at RWTH Aachen University.

## Author Contributions

Conceptualization: S.E., and A.S.; methodology: S.E., L.G., S.H., and A.Se.; software: S.E., S.S., S.H., N.T.; investigation: S.E., L.G., S.H., T.R.; resources: N.W., F.K., T.C.; funding acquisition: A.S., N.W., F.K., and T.C.; supervision: A.S. and N.W.; original draft preparation: A.S., S.E., N.T., N.W., and T.C.; review and editing: A.Se., L.G., S.S., S.H., T.R., and F.K.

## Data availability statement

The microbiota sequencing data presented in this study were submitted to the Sequence Read Archive and are openly available under the accession number PRJEB80543.

## Supplementary information

The online version contains supplementary material available at https://doi.org/xx/scientificreports.

