## Supplemental Information for "Intercellular adhesion molecule-1 protects against adipose tissue inflammation and insulin resistance but promotes liver inflammation and hepatic fibrosis in mice"

### Supplementary Figures:

#### Supplementary Figure S1:

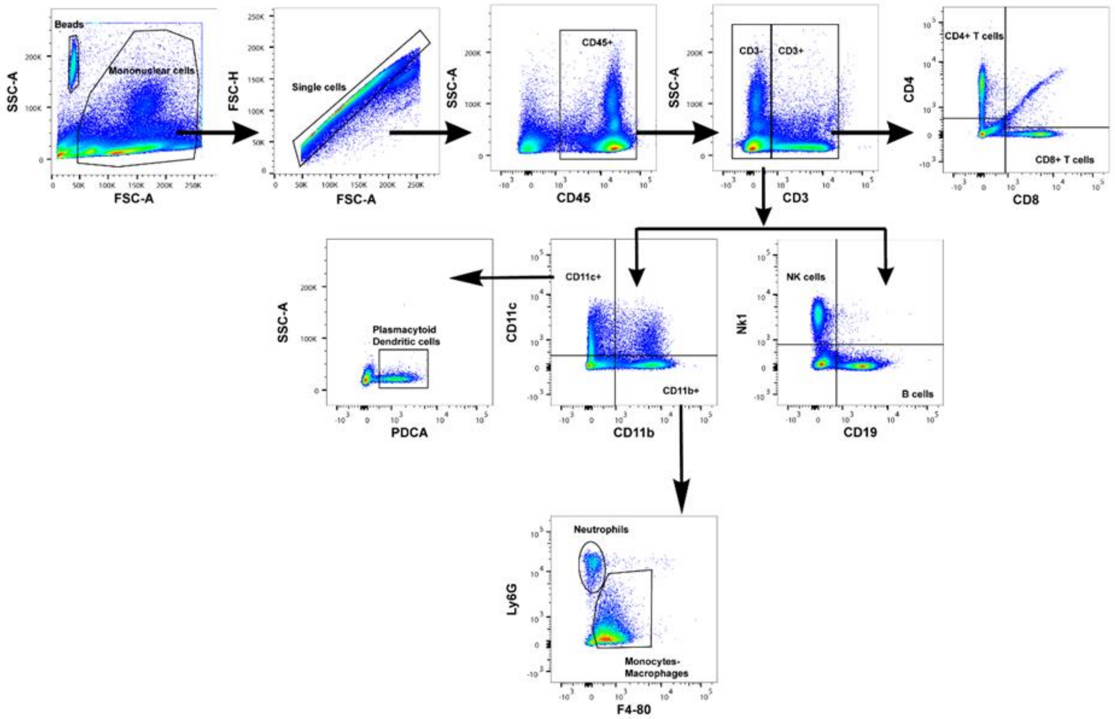


**Supplementary Figure S1.** Immune cell subset gating by multiparameter flow cytometry (liver cells, representative of all organ fractions). The analysis included: B cells (CD45$^+$CD3$^-$CD19$^+$), CD4$^+$ T cells (CD45$^+$CD3$^+$CD4$^+$), CD8$^+$ T cells (CD45$^+$CD3$^+$CD8$^+$), natural killer (NK) cells (CD45$^+$ CD3$^-$NK1.1$^+$), monocytes/macrophages (Mo-MF)(CD45$^+$CD11b$^+$Ly6G$^-$ F4/80$^+$), neutrophils (CD45$^+$CD11b$^+$ CD11c$^-$F4/80$^-$Ly6G$^+$), and plasmacytoid dendritic cells (pDCs) (CD45$^+$ CD11b$^+$CD11c$^+$PDCA$^+$).

**Supplementary Figure S2:**

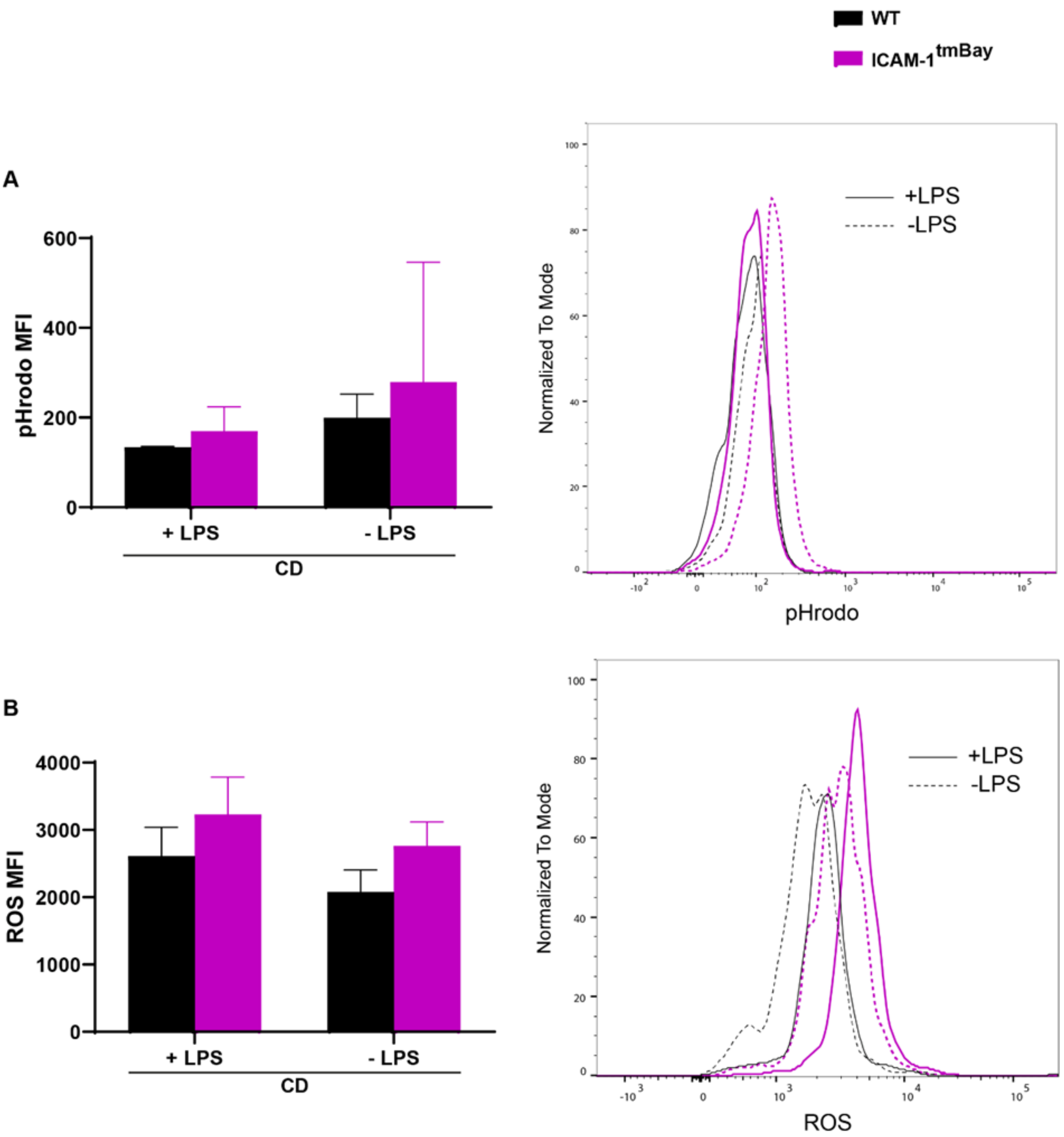


**Supplementary Figure S2.** Flow cytometric analysis of pHrodo and reactive oxygen species (ROS) in LPS-stimulated or unstimulated $CD45^+CD11b^+Gr1^+$ myeloid cells of whole blood from chow-fed (CD) WT mice (shown in black, n = 3) and ICAM-1 mutant ($Icam1^{tmBay}$) mice (shown in purple, n = 5). (A) Mean fluorescence intensity (MFI) of pHrodo and representative histograms. (B) Mean fluorescence intensity (MFI) of ROS and representative histograms. Statistical significance was calculated by two-way ANOVA (A-B). Values are represented as mean ± SD.

**Supplementary Figure S3:**

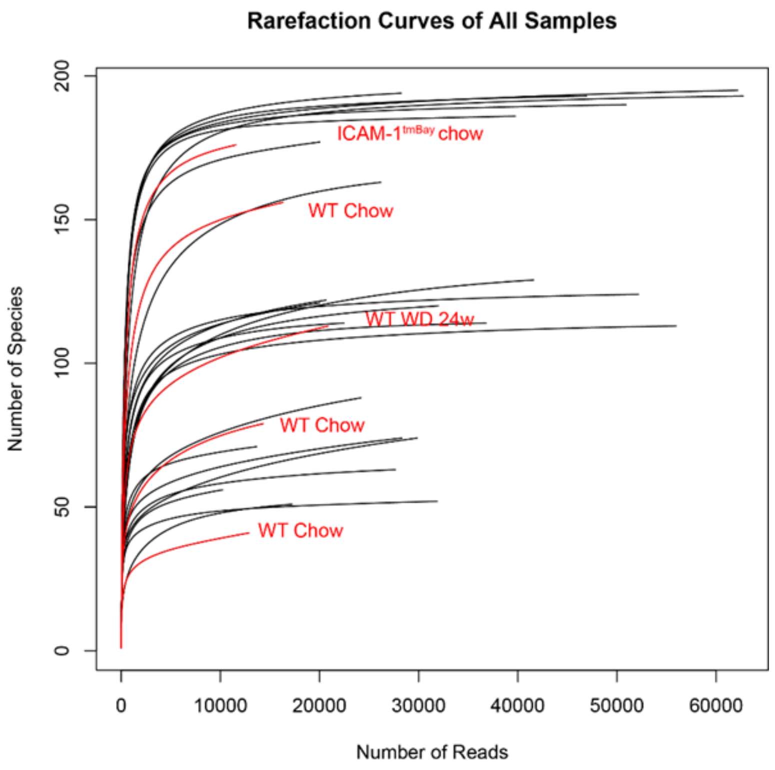


**Supplementary Figure S3.** Rarefaction curves depicting sequencing depth. The number of observed molecular species in each sample was plotted against the number of reads acquired, to ensure sufficient sequencing depth. The curve was generated in R using Rhea [1]. The five samples with the least sequencing reads are shown in red.

**Supplementary Tables:**

**Supplementary Table S1:** Antibodies for histochemical analysis

| Antibody | Channel/Conjugate | Manufacturer | Clone | Dilution |
|---|---|---|---|---|
| **ICAM-1** | Pure | Invitrogen | 020 | 1:50 |
| **MPO** | Pure | Bioss | bs4943R | 1:100 |
| **CD68** | Pure | Abcam | ab125212 | 1:200 |
| **Goat anti-rabbit IgG** | Biotin | VectorLabs | BA-1000 | 1:200 |

ICAM-1, Intercellular adhesion molecule 1; MPO, Myeloperoxidase.

**Supplementary Table S2:** Antibodies used for flow cytometric analysis

| Antibody | Channel/Conjugate | Manufacturer | Clone | Dilution |
|---|---|---|---|---|
| **NK 1.1** | AF 488 | Biolegend | PK136 | 1:400 |
| **CD4** | PE | BD biosciences | GK1.5 | 1:400 |
| **CD45** | APC-Cy7 | BD biosciences | 30-F11 | 1:400 |
| **CD19** | efluor 450 | eBioscience | 1D3 | 1:200 |
| **CD8a** | Amcyan | Biolegend | 53-6.7 | 1:400 |
| **CD3e** | PerCp-Cy5.5 | eBioscience | 145-2C11 | 1:100 |
| **F4/80** | efluor450 | Serotec | CI:A3-1 | 1:200 |
| **CD11b** | Amcyan | BD Biosciences | M1/70 | 1:400 |
| **Ly6G** | FITC | BD Biosciences | 1A8 | 1:400 |
| **CD317** | PE | eBioscience | eBio129c | 1:400 |

AF 488, Alexa Fluor 488; PE Phycoerythrin; APC-Cy7, Allophycocyanin-Cyanine 7; FITC Fluorescein isothiocyanate.

**Supplementary Table S3:** Primers used in this study

| Primer | Forward | Reverse |
|---|---|---|
| ***Gapdh*** | acctgccaagtatgatgacatca | ggtcctcagtgtagcccaagat |
| ***Icam-1*** | cacccaccccgcaggtcca | ttccccaagcagtccgtctcg |
| ***Mcp-1*** | agagccagacgggaggaag | ccagcctactcattgggatc |
| ***Col1a*** | gcagggttccaacgatgttg | gcagccatcgactaggacaga |

*Gapdh*, Glyceraldehyde 3-phosphate dehydrogenase; *Icam-1*, Intercellular adhesion molecule 1; *Mcp-1*, Monocyte chemoattractant protein-1; *Coll1a*, Collagen type I alpha.
